# TISSLET Tissues-based Learning Estimation for Transcriptomics

**DOI:** 10.1101/2025.02.18.638832

**Authors:** Ahmed Miloudi, Aisha Al-Qahtani, Thamanna Hashir, Mohamed Chikri, Halima Bensmail

## Abstract

In the context of multi-omics data analytics for various diseases, transcriptome-wide association studies leveraging genetically predicted gene expression hold promise for identifying novel regions linked to complex traits. However, existing methods for multi-tissue gene expression prediction often fail to account for tissue-tissue expression interactions, limiting their accuracy and effectiveness.

This research addresses the challenge of predicting gene expression across multiple tissues by incorporating tissue-tissue expression correlations based on a nonlinear multivariate model. Our findings demonstrate that this model excels in estimating tissue-tissue interactions and accurately predicting missing data. These results have significant implications for multi-omics data analytics and transcriptome-wide association studies, suggesting a novel approach for identifying regions associated with complex traits.

## 1 Introduction

eQTL (expression Quantitative Trait Loci) studies explore the relationship between genetic variants, such as SNPs (Single Nucleotide Polymorphisms), and gene expression levels. By identifying how specific SNPs influence gene expression, researchers can gain insights into the genetic basis of complex traits and diseases. In the context of transcriptomics, which involves the comprehensive analysis of RNA transcripts, eQTL mapping is particularly valuable (Figure 1). When considering multiple tissues, incorporating tissue-tissue correlations becomes essential, as gene expression can be influenced by interactions across different tissues. This approach allows for a more holistic understanding of gene regulation and the identification of novel regions associated with complex traits, enhancing the precision and effectiveness of transcriptome-wide association studies. Transcriptome-wide association studies (TWAS) have therefore gained prominence in the field of genomics and genetics for their potential to uncover genetic variants associated with complex traits. These studies leverage gene expression data to bridge the gap between genetic variations and phenotypic traits. Although TWAS holds great promise, the accuracy of gene expression prediction across multiple tissues remains a challenge. Previous research addressed the challenge of multi-tissue gene expression (Grinberg and Wallace, 2021) [1]

**Fig. 1.**
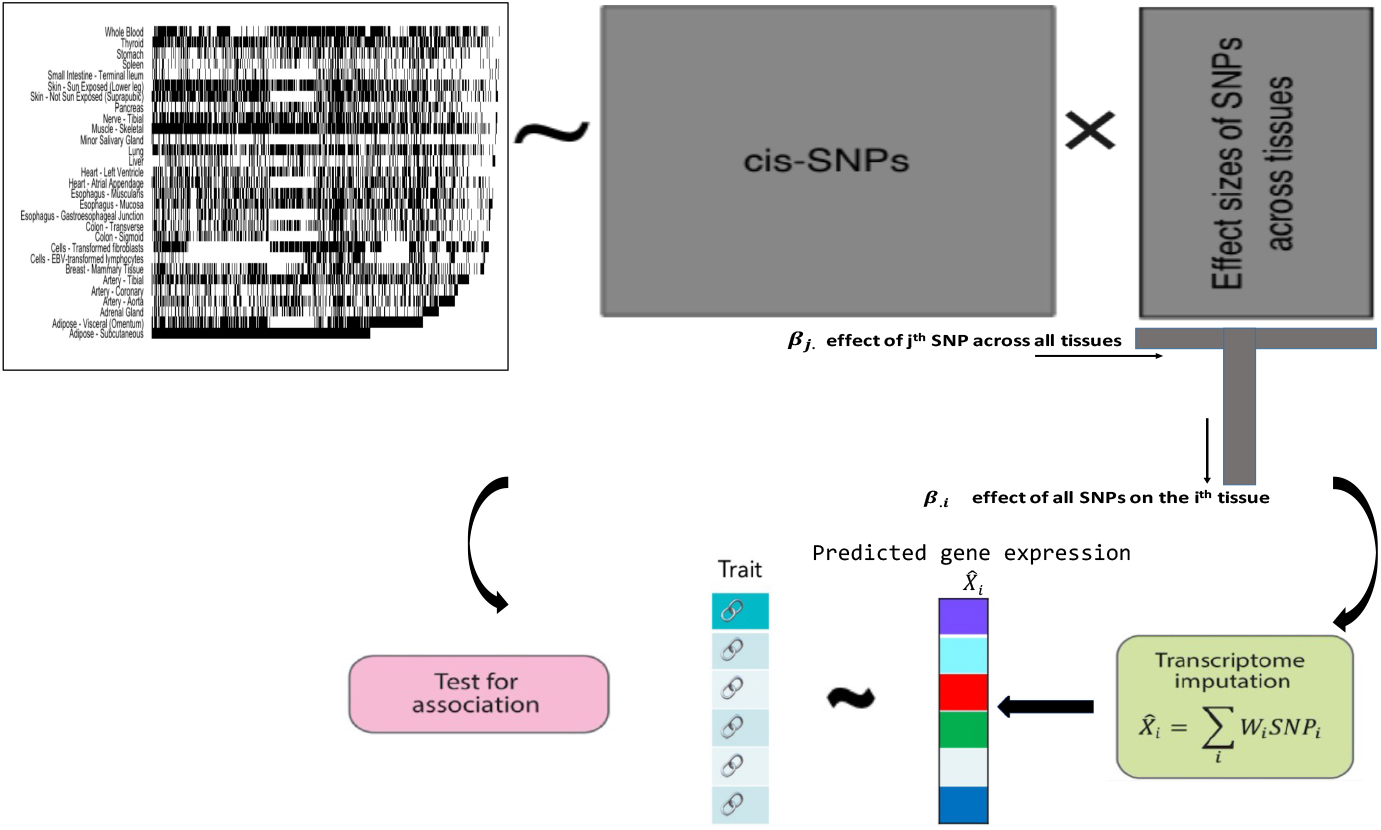
Overview of TISSLET method. The inputs required for TISSLET is a matrix of gene expression for several tissues. We also use the subjects genotypes with sample matched measured expression. TISSLET’s weights output are calculated based on the CEM algorithm using measured gene expression and provide the weights and covariance structure of tissues. For full details, see the Materials and Methods section.

Machine learning algorithms have been used to predict eQTL regulation of a gene by analyzing adjacent genetic variants [2, 3]. Researchers have used genetically predicted gene expression models to conduct transcriptome-wide association studies (TWAS) and identify novel areas linked to complex traits. However, many of these regions lack a GWAS relationship within 1Mb.

There are various benefits to such analyses: Leveraging gene expression enriches possible trait-associated SNPs, whereas joint eQTL modeling improves overall association strength and reduces the number of tests from millions to roughly 20,000 genes.

Leveraging shared eQTLs across tissues enhances eQTL discovery and gene expression imputation accuracy, leading to more powerful transcriptome-wide association analyses [4–6]. Hu et al. [7] and Molstad et al. [8] developed a penalized regression strategy for collaborative modeling of eQTLs. The penalty encourages shared eQTLs between tissues. Using genotype and expression data from the Genotype-Tissue Expression (GTEx) project, multi-tissue eQTL models significantly increase gene association identification and imputation accuracy compared to single-tissue techniques.

While Hu et al.’s technique [7] does not account for tissue-specific gene expression correlation in combined eQTL modeling, Molstad et al. [8] uses cross-tissue imputation with EM algorithm. Both approach uses linear model to derive their algorithm. Recent research suggests that accounting for metric based on skewness improves variable selection and prediction accuracy and can identify regulators or genes in large patient cohorts. A recent study showed that significant correlation was detected between expression skewness and the top 500 genes corresponding to the most significant differential DNA methylation occurring in the promotor regions in TCGA [9]

Recent research suggests that accounting for tissue-tissue correlation in high-dimensional penalized multivariate response linear regression improves variable selection and prediction accuracy. This phenomena may be explained by a seemingly unrelated regression interpretation of high-dimensional sparse multivariate response linear regression. Moreover, certain tissue types are more difficult to obtain due to biological and cost constraints. Using tissue-tissue correlation with skewness assumption enhances gene expression prediction accuracy, particularly for small data numbers.

In this paper, we propose an approach (TISSLET) which not only impute missing gene expression using cross information from several tissue, but also estimate tissue-tissue correlation. We calculate a joint eQTL weights while imputing missing gene expression using a skewed normal modeling. The technique by which our approach works is straightforward. Measuring expression in one tissue can provide a reliable estimate of expression in the other, especially if their expression is substantially correlated. Ignoring tissue-tissue correlation can significantly reduce gene expression prediction accuracy. Our methodology offers several advantages over previous approaches for multi-tissue joint eQTL mapping, including:

1. Incorporation of a full tissue-tissue correlation matrix in the model, rather than assuming a diagonal matrix, which reveals cross-tissue expression dependencies that eQTLs alone cannot explain.
2. Efficient estimation of eQTL weights by modeling cross-tissue associations.
3. Relaxation of the normality assumption by allowing for a heavy-tailed error distribution, assuming that errors follow a multivariate skewed distribution.

Figure 1 gives an overview of TISSLET method. This figure demonstrates the training of the imputation model. The inputs required for TISSLET is a matrix of gene expression for several tissues. We also use the subjects genotypes with sample matched measured expression. TISSLET’s weights output are calculated based on the CEM algorithm using measured gene expression and provide the weights and covariance structure of tissues. For full details, see the Materials and Methods section.

## 2 Materials and Methods

Regression models are commonly used to map the relationship between SNPs and gene expression levels in eQTL studies. Traditional models often assume normality of errors, which may not capture the true distribution of gene expression data. Introducing a skewed model with a cross-tissue expression based on SNP genotypes can provide a more accurate representation by allowing for heavy-tailed error distributions, improving the precision of eQTL mapping and the identification of genetic influences on gene expression.

Let **x**_*i*_ ∈ℝ^*p*^ represent the genotypes of *p* SNPs (both centered and normalized) and let **y**_*i*_ ∈ℝ^*q*^ represent the vector of centered and normalized measured expression in *q* tissues for the *i*^*th*^ subject for a specific gene within a certain distance (e.g. 500 kb) or less away from the gene of interest. We assume that gene expression represents a realization of the random vector for the *i*^*th*^ subject:

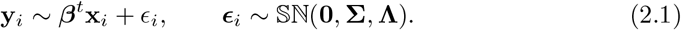

where 𝕊ℕ_*q*_ denotes the q-dimensional multivariate skewed normal distribution, where ***β*** ∈ ℝ^*p×q*^ is the unknown regression coefficient matrix (i.e., eQTL weights), and **S**^−**1**^ ∈ ℝ^*q×q*^ is the cross-tissue error precision (inverse covariance) matrix. We further assume that *ϵ*_*i*_ is independent of *ϵ*_*j*_ for all *i* ≠ *j*.

The skewed distribution 𝕊ℕ can reveal the asymmetric information when the observations such as gene expression are skewed [10]. Asymmetrical distributions have an additional shape parameter **Λ** ∈ℝ^*q×q*^ to represent the direction of the asymmetry of the density. If the skewness in observations is ignored, inferences with symmetric distributions may result in biased or even misleading conclusions.

### 2.1 Penalized skew-normal log-likelihood

In appendix (7.2) we outline in details the derivation of the model parameters (weights and cross-correlation matrices) under the normality assumption. This approach uses the ECM algorithm for parameter estimation. Algorithm 1 gives the pseudocode of the approach using normality assumption.

#### Algorithm 1

Regularized ECM Algorithm with normality

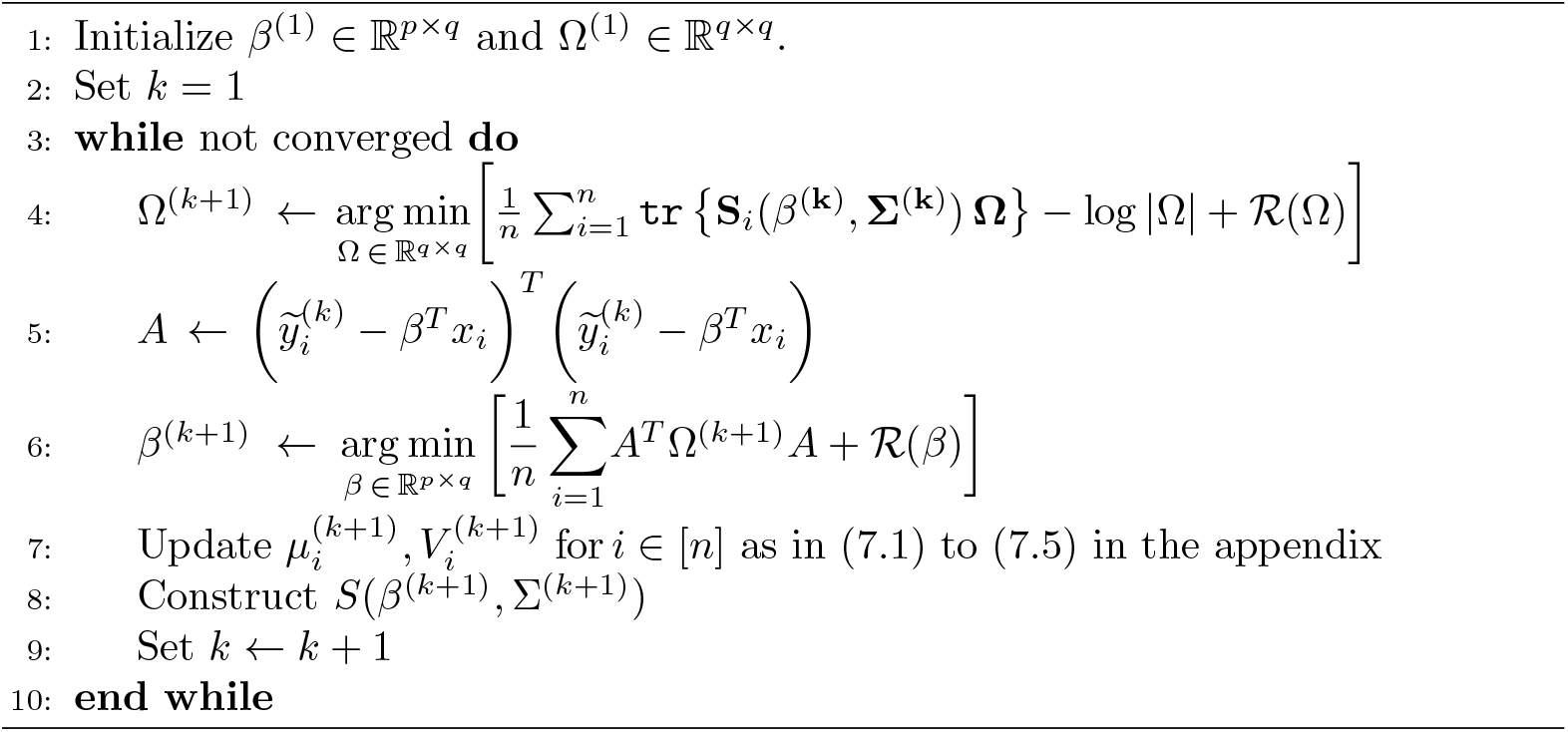

Similarly, using equation (2.1) and assuming that the errors *ϵ*’s 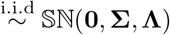,for 1 ≤*i* ≤*n*, and assuming that gene expression represents a realization of the random vector for the *i*^*th*^ subject:

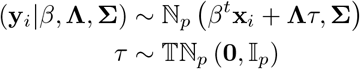

where **x**_*i*_ = (**x**_*i*,1_, …, **x**_*i,p*_) is the vector of covariates (here **x**_*i*_ is the *i*^*th*^ individus genotype), *β*^*t*^ = (*β*_1_, …, *β*_*p*_)^*t*^, is an unknown matrix of mean regression coefficient and 𝕋ℕ is a truncated normal distribution.

On the other hand, since gene expression matrix has random missing values, we consider that 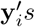 are partially observed with an arbitrary missing pattern. In order to set up estimating equations for multivariate data with possible missing values, we separate **y**_*i*_(*q ×* 1) into two components 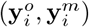 accordingly, where 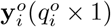 is the observed c mponent and 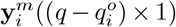 is the missing component. Further, we introduce two missingness indicator matrices, denoted by **O**_*i*_ and **M**_*i*_ henceforth, corresponding to **y**_*i*_ such that 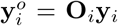 and 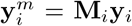,respectively. More specifically, 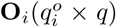 and 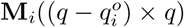 are sub-matrices extracted from the rows of an identity matrix of order *q*, **I**_*q*_, corresponding to row positions of 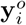 and 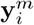 in **y**_*i*_, respectively. When **y**_*i*_ is fully observed, **O**_*i*_ = **I**_*q*_ and **M**_*i*_ is null. Meanwhile, it is easy to verify that: 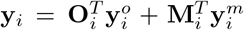 and 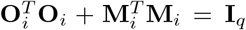.Using appendix (7.3) and Bayes Theorem the negative log-likelihood for the observations **y**_1_, **y**_2_, …, **y**_*n*_

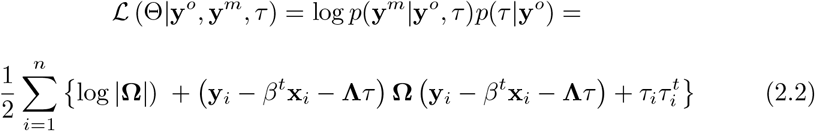

Then the expected conditional log likelihood **Q**(*β*, **Ω, Λ**) is expressed as

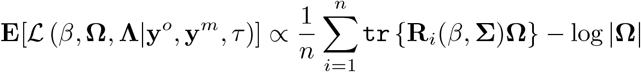

where

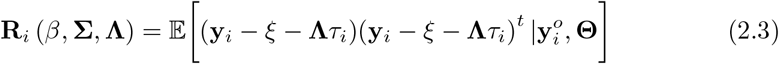

### 2.2 Regularization for precision matrix Ω

We regularize the entries of **Ω** using *𝓁*_1_ penalties, then the regularized conditional log likelihood **Q** is expressed as:

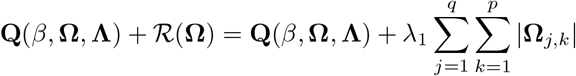

For sufficiently large tuning parameter values *λ*_1_, the penalty results in precision matrix estimations **Ω** with all off-diagonal entries equal to zero. The penalty assumes that some entries are equal to zero. In multi-tissue joint eQTL mapping, a zero in the (j,k)^*th*^ entry of **Ω** indicates independent expression in the j^*th*^ and k^*th*^ tissues, given expression in all other tissues and p SNP genotypes (1). We do this for two reasons: first, when *p > n* (which is the case for practically every gene we test, more SNPs than observations), without penalizing the diagonals, a perfect fit can occur in the M-step and not in E step (see Algorithm 2): Recent literature, has shown that precision matrix estimators employed in predictive models exhibit this behavior [11]. Next we summarize the main ECM Algorithm 2. Algorithm 3 gives a detailed version of Algorithm in appendix 7.3):

#### Algorithm 2

Regularized ECM Algorithm with non-normality

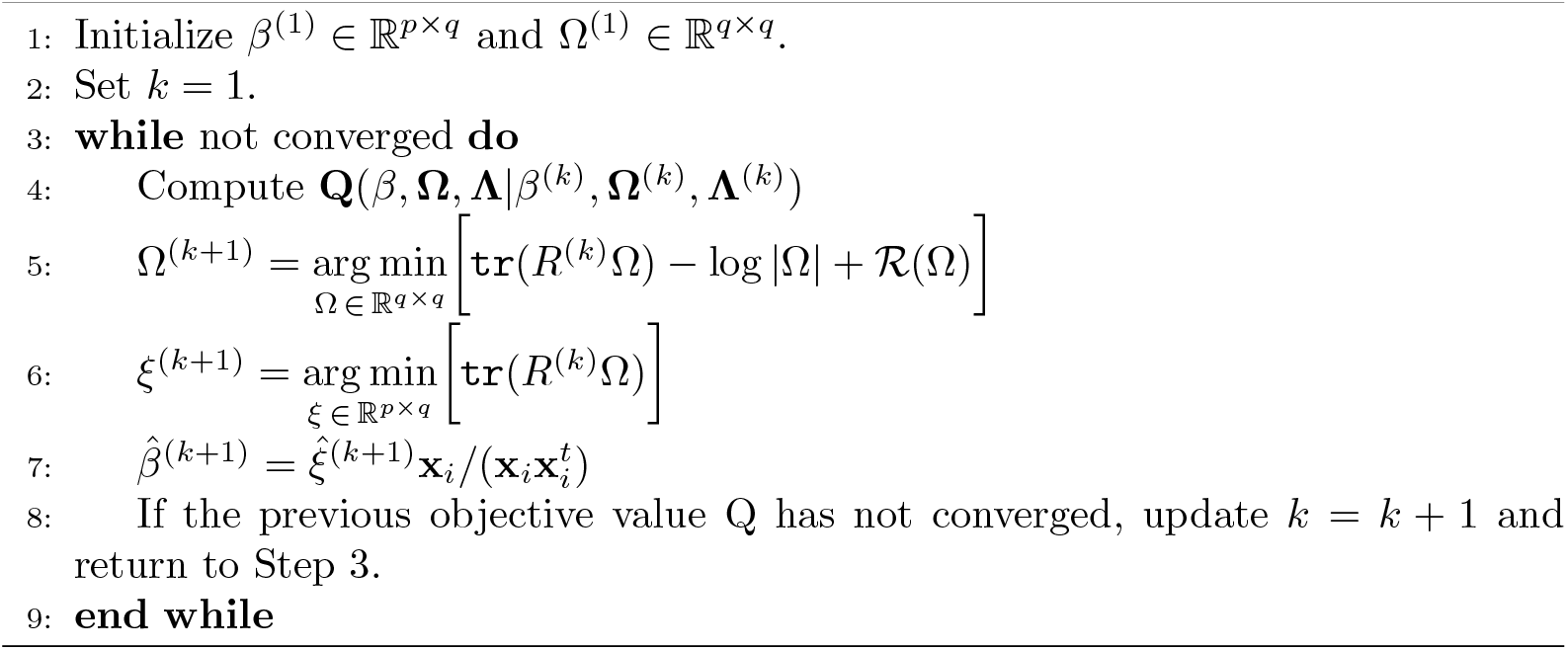

#### Remark 1

**Step 4 of Algorithm 2 can be expressed**

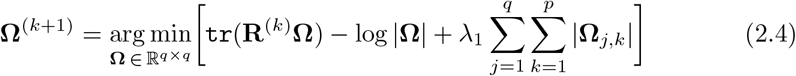

Conveniently, (2.4) is exactly the optimization problem for computing the *𝓁*_1_-penalized normal log-likelihood precision matrix estimator with input sample covariance matrix **R**(.). Many efficient algorithms and software packages exist for computing (2.4). We can use our R software BiGQUIC to solve (2.4) which is available at BiGQUIC [12]

#### Remark 2.

**Step 5 of Algorithm 2 can be expressed as:**

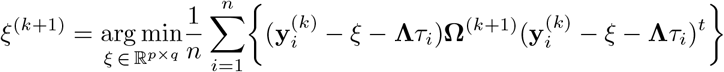

Using appendix (7.3), we have

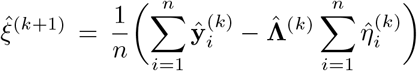

where

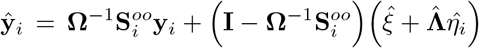

## 3 Illustrations

### 3.1 Simulation study

In this section, we utilize 80 replications of data generated from multivariate regression, where the sample size is *n* = 50, 150, *p* = 22, and *q* = 24. The choice of *q* is made to align with the dimension of the regression models applied to the GTEx data. In each replication, a sparse matrix **B** is generated using the element-wise product of three matrices: **B** = **W** ⊙ **K** ⊙ **Q** where ⊙ is the elementwise product, (**W**)_*ij*_ ∼ *N* (0, 1) and (**K**)_*ij*_ ∼ *Bernouilli*(*s*_1_) and each row of **Q** consists of either all 1’s or all 0’s with a success probability of 1’s equal to *s*_2_.

By generating **B** in this manner, we anticipate that (1 − *s*_2_)*p* predictors will be irrelevant for all *q* responses, and each predictor will be relevant for *s*_1_*q* of all the response variables. An *n × p* predictor matrix **X** is also generated with rows drawn independently from *N* (**0, ∑**_**X**_), where (**∑**_**X**_)_*ij*_ = 0.7^|*i*−*j*|^, following the approach of Yuan and Lin (2007) [13] and Peng et al. (2010) [14]. We consider an AR(1) covariance structure for the scale matrix of the errors, which is ∑ = *ρ*^|*i*−*j*|^.

Lastly, every row of the error matrix **E** is independently sampled from a multivariate skew-Normal distribution 𝕊 ℕ (**0, ∑, Λ**), and the response matrix **Y** is formed as **Y** = **XB** + **E**. To reduce computation time, we independently generate validation data (sample size *n* = 50) within each replication to estimate the prediction error for the algorithms, akin to performing a K-fold cross-validation for the algorithm, as described in Rothman et al. (2010) [15].

We consider 36 different combinations of **Λ**; *ρ*; *s*_1_ and *s*_2_ from the following ranges:(1) 7 *ρ* = {0, 0.4, 0.8}, (2) **Λ** = *diag*(−1, 1, −1, …, 1) or 𝟙_*q*_, where 𝟙_*q*_ is a column vector of ones, (3) *s*_1_ = 0.1, 0.5, and (4) *s*_2_ = 1. Tuning parameters is selected from the set {2^*a*^, *a* ∈ 0; *±*1, …, *±*5}.

### 3.2 Results on simulation study

In the simulation study, we measure the overall performance of various methods in terms of the mean squared prediction error (PE). We have computed the prediction errors (PE) values of the entropy loss functions of the estimators of **Ω** for the 80 simulated datasets using the k-NN, Normal, the G-lasso and the TISSLET algorithm (Figure 2). To assess the significance of the difference in prediction error between the Normal and Tisslet algorithms, we conducted a Wilcoxon signed-rank test (*R*_*i*_ = *Sign*(*D*_*i*_) −*rank*(|*D*_*i*_|) where *D*_*i*_ is the difference of both predictions. The results indicated a statistically significant difference, with a p-value of 0.0124 at the 5% significance level (*α* = 0.05).

**Fig. 2.**
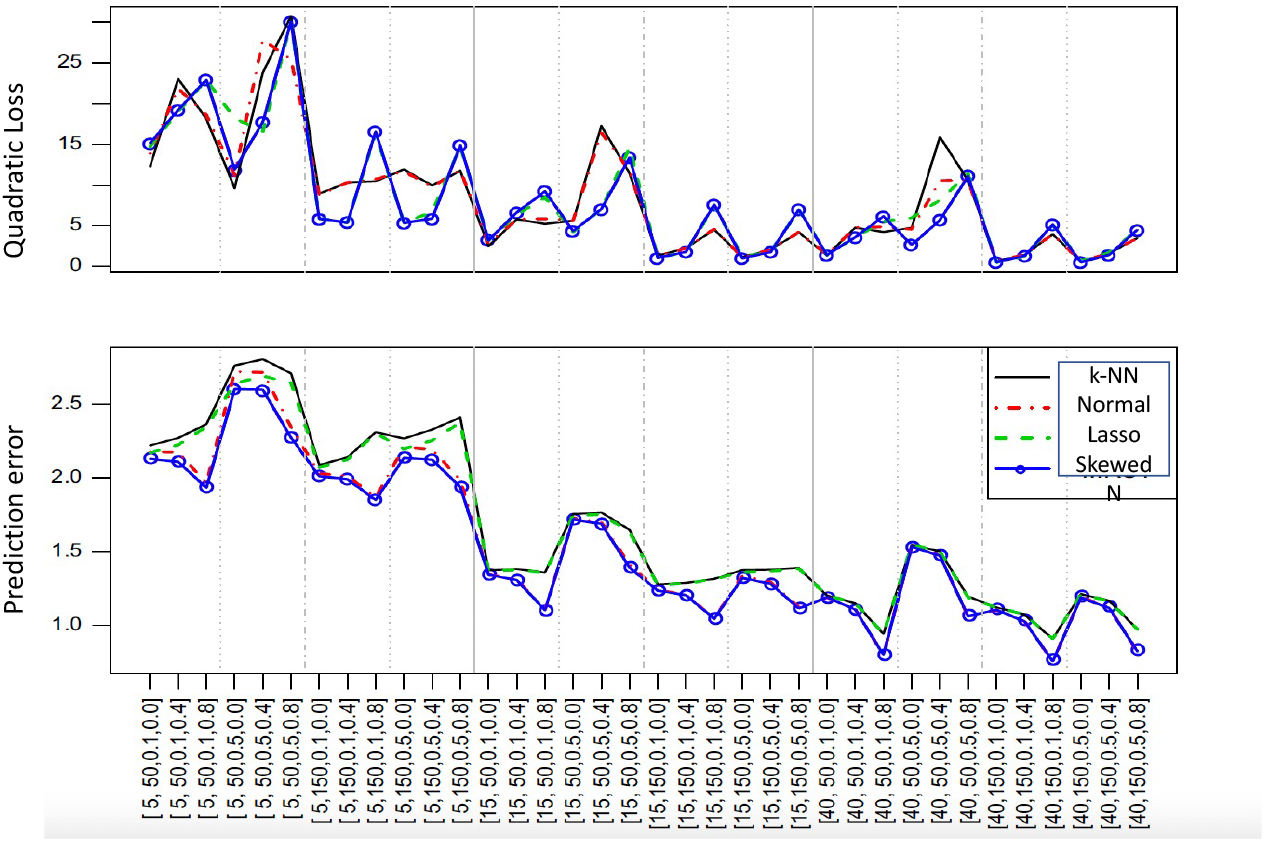
Prediction errors (PE) values of the entropy loss functions of the estimators of **Ω** for the 80 simulated datasets using the k-NN, Normal, the G-Lasso and the TISSLET (skewed) algorithm

While the *𝓁*_1_ loss is used to evaluate the performance of **Ω**, the sparsity recognition performance of **Ω** is measured by the true positive rate (TPR) as well as the true negative rate (TNR) defined as

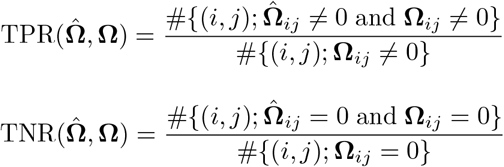

The corresponding true positive rates (TPR) and true negative rates (TNR) for are reported in Tables 1 and 2. From these tables it is evident that a slightly higher TPR is accompanied by a lower TNR for our algorithm Tisslet algorithm versus Molstad et al. algorithm (normal assumption). We have also compared the numerical performance of the algorithms and the G-Lasso method obtained from Friedman et al. (2022) [16] by setting Λ to be **0** for the 36 combinations. In general, it turns out that for higher error correlations (*ρ* = 0.8) the mean square error of these methods is somewhat lower compared to the K-NN method, but the estimators of **Ω** are improved considerably. Finally, the computational times of the two algorithms are computed. The computational times for the four algorithms were evaluated and compared. On average, the CPU time ratios of Normal, K-NN, and Lasso to our algorithm, Tisslet, were 3.79, 4.72 and 2.17 respectively. These computations were performed on the Panther cluster, equipped with 2.5 GHz processors. The differences in computational time are attributed to their computational complexities, which are generally *O*(*np*), for all except for K-NN, which has a complexity of *O*(*kn*^2^*p*). Lasso exhibits a lower ratio compared to the other algorithms because it penalizes small values in the covariance matrix, effectively reducing the time spent on its computation.

**Table 1.**
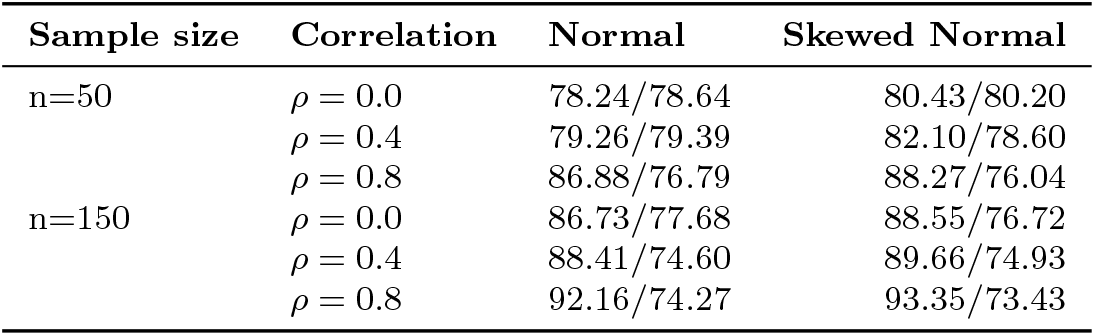
TPR/TNR for the matrix **Ω** averaged over 80 replications with *s*_1_ = 0.1, *s*_2_ = 1 and λ = (1; 1; 1; … ; 1)^*T*^.

| Sample size | Correlation | Normal | Skewed Normal |
| --- | --- | --- | --- |
| n=50 | $\rho = 0.0$ | 78.24/78.64 | 80.43/80.20 |
| | $\rho = 0.4$ | 79.26/79.39 | 82.10/78.60 |
| | $\rho = 0.8$ | 86.88/76.79 | 88.27/76.04 |
| n=150 | $\rho = 0.0$ | 86.73/77.68 | 88.55/76.72 |
| | $\rho = 0.4$ | 88.41/74.60 | 89.66/74.93 |
| | $\rho = 0.8$ | 92.16/74.27 | 93.35/73.43 |

**Table 2.**
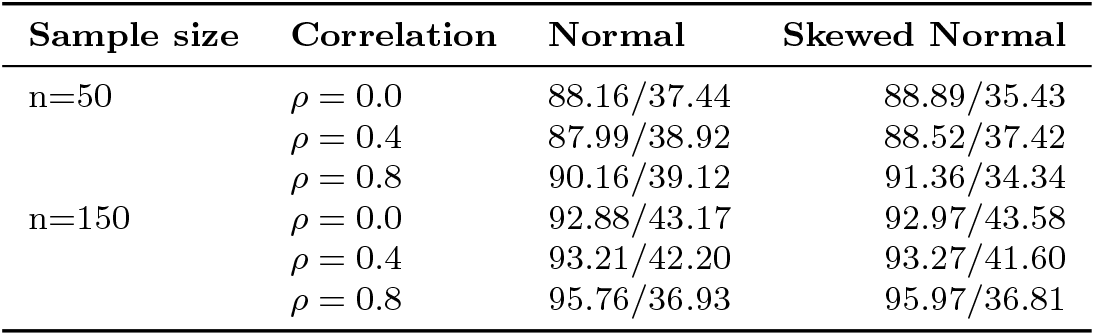
TPR/TNR for the matrix **Ω** averaged over 80 replications with *s*_1_ = 0.5, *s*_2_ = 1 and λ = (1; 1; 1; … ; 1)^*T*^.

### 3.3 Data generating study for SPATC1L gene

We conducted extensive numerical experiments to examine how the number of shared eQTLs, population R^2^ (also known as narrow-sense heritability), and how tissue-tissue correlation structure affect the performance of various methods for estimating eQTL weights across multiple tissues. To closely replicate the conditions of joint eQTL mapping in GTEx data, we obtained whole-genome sequencing SNP genotype data for all SNPs within 500kb of the SPATC1L gene for 620 subjects from the GTEx dataset [17]. After removing highly correlated SNPs (see Data Preparation section), we were left with *p* = 1178 SNP genotypes. For each replication, we then generated *n* = 620 subjects’ expressions in *q* = 29 tissues. Denoting **x**_*i*_ ∈ ***R***^*p*^ as the SNP genotypes for the i^*th*^ subject, we generated **y**_*i*_ ∈ *ℝ*^*q*^ as a realization of the random vector 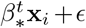 for *i* = [*n*], where *β* ∈ *ℝ*^*p×q*^ are the eQTL weights and errors are independent and identically distributed as 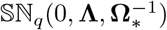 (Same protocol as [8] and [7]). For five hundred independent replications in each scenario, we randomly divided the *n* = 620 subjects into a training set of size *n*_train_ = 400, a validation set of size 110, and a testing set of size 110.

To create missingness, in each independent replication, we generated **y** as follows: first, we created a matrix **B** ∈ ℝ^*p×q*^ with entries that were independent 𝒩 (0, 1). Then, we generated a matrix **S** ∈ ℝ^*p×q*^ to be a matrix whose rows are either all zero or all one: we randomly selected *s* rows to be nonzero, where *s* ∈ [20]. Given **S**, we then generated a matrix **U** ∈ ℝ^*p×q*^ so that each of the *q* columns has 20 − *s* randomly selected entries equal to one only from entries which are zero in *S* and all others equal to zero.

We calculated **y** = **B** ⊙**S** + **B** ⊙ **U**, where ⊙ denotes the element-wise product. This construction ensured that each tissue has twenty total eQTLs, *s* of which are shared across all tissue types. We considered *s* = {5, 10, 15, 18, 20} in the simulations presented in this section. It is important to note that due to high linkage disequilibrium, many SNPs are highly correlated, resulting in a larger number of SNPs associated with gene expression. Figure 3 gives a summary of the protocol for generating data. In addition, we randomly introduced missing values to both the training and validation set responses with a missing probability of 0.55, which corresponds to the missing rate in the GTEx gene expression data. For each method, we trained the model on the training data, chose tuning parameters using the validation data, and evaluated prediction and variable selection accuracy on the testing data.

**Fig. 3.**
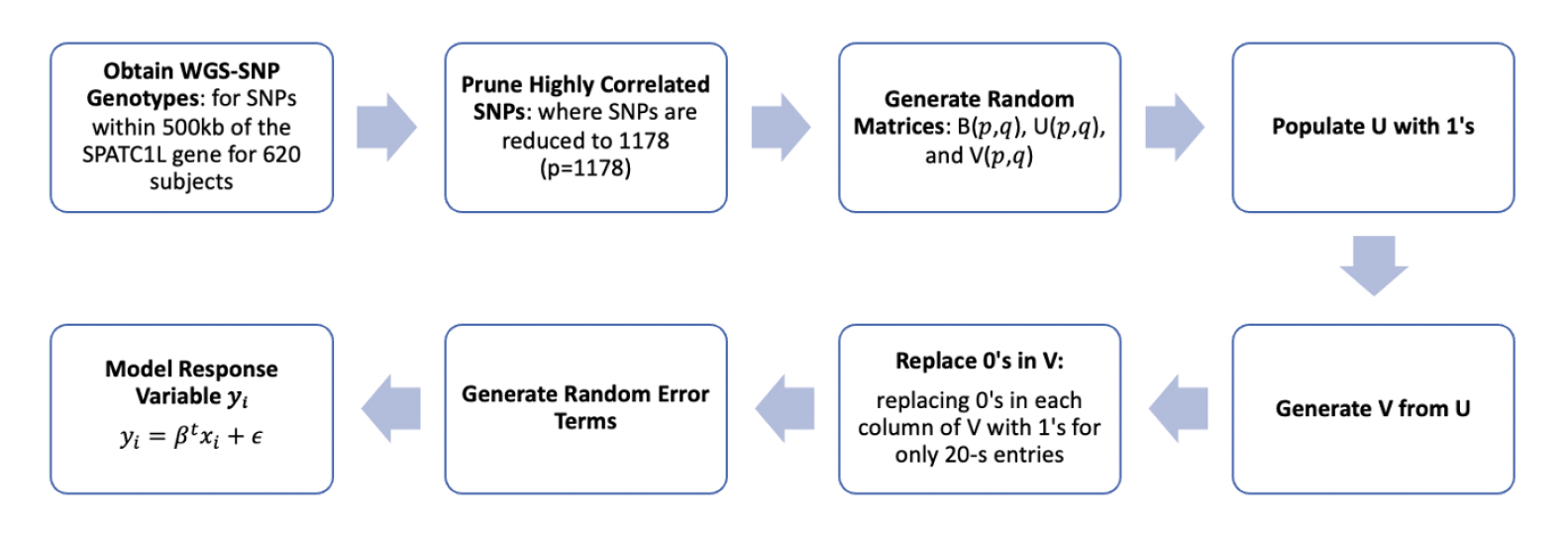
Overview of the Data Pre-processing Steps. Input WGS SNP data then prune highly correlated SNPs. Create U and V and generate B. Generate error annd fit the model.

To construct the covariance matrix **Ω**, we build it to have a block-diagonal structure and to control the R^2^. Specifically, we consider a covariance structure for the scale matrix of the errors, which is **S**_*ij*_ = *ρ*^|*i*−*j*|^, for *i* ∈ [20, 20] and **S**_*ij*_ = (*ρ* + 0.2)^|*i*−*j*|^, for *i* ∈ [10, 10].

### 3.4 Results on SPATC1L gene study

#### Comparative Analysis of Covariance Approaches

We evaluated the *s*-TISSLET imputation method’s performance at different values of *s* for imputing missing data generated in section (3.3). Figure 4 shows that despite the varying *s* values, the imputed distributions closely align with the original distribution. This indicates that the imputation method maintains the overall structure of the gene expression data, which is critical for the reliability of subsequent analyses.

**Fig. 4.**
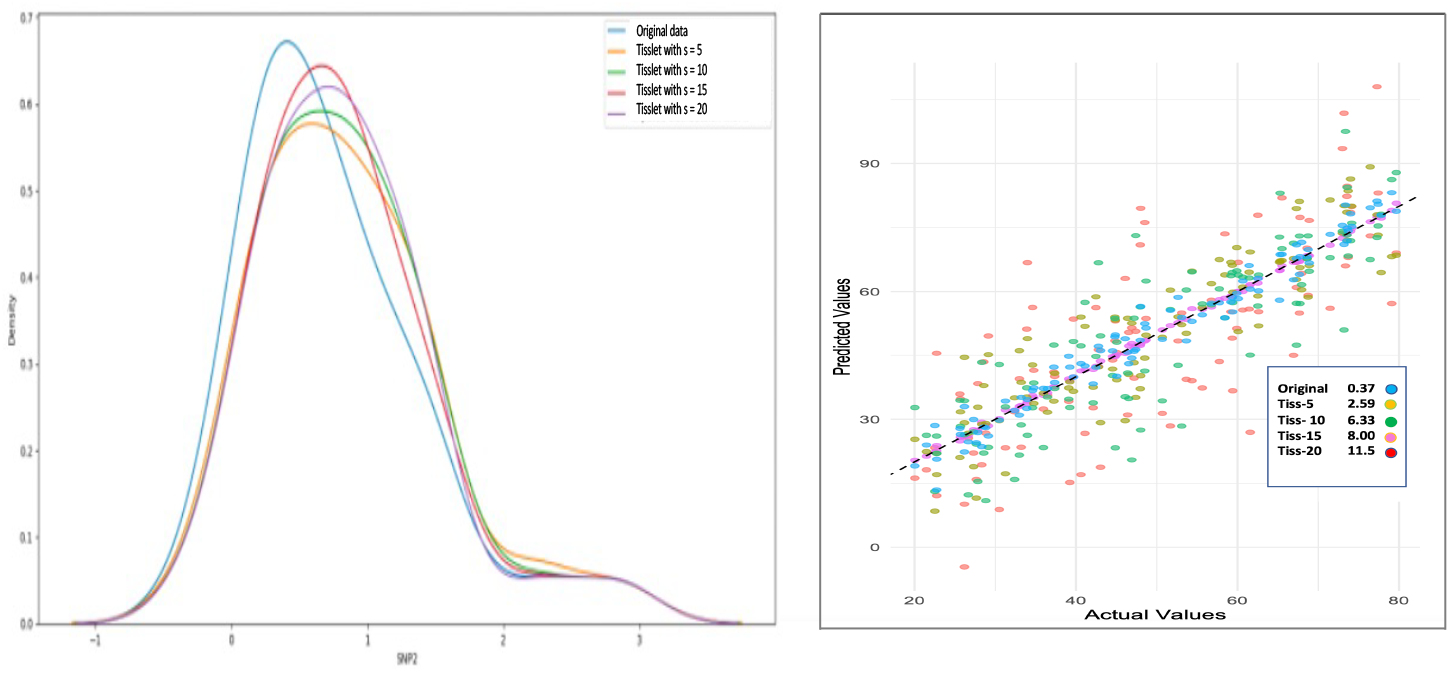
Left: comparing distribution of original versus Tisslet imputed SNP_2_ using several choice of *s* (5, 10, 15, 20) distributions. Right: providing the Mean absolute error (MAE)

#### Dataset size and prediction accuracy

In the assessment of the TISSLET framework’s performance, a nuanced relationship between dataset size (*N*) and R^2^ scores became apparent (Figure 5a). An increase in *N* was correlated with a decline in R^2^ scores, underscoring the challenges in sustaining prediction accuracy with the expansion of data volume inherent in multi-omics studies.

**Fig. 5.**
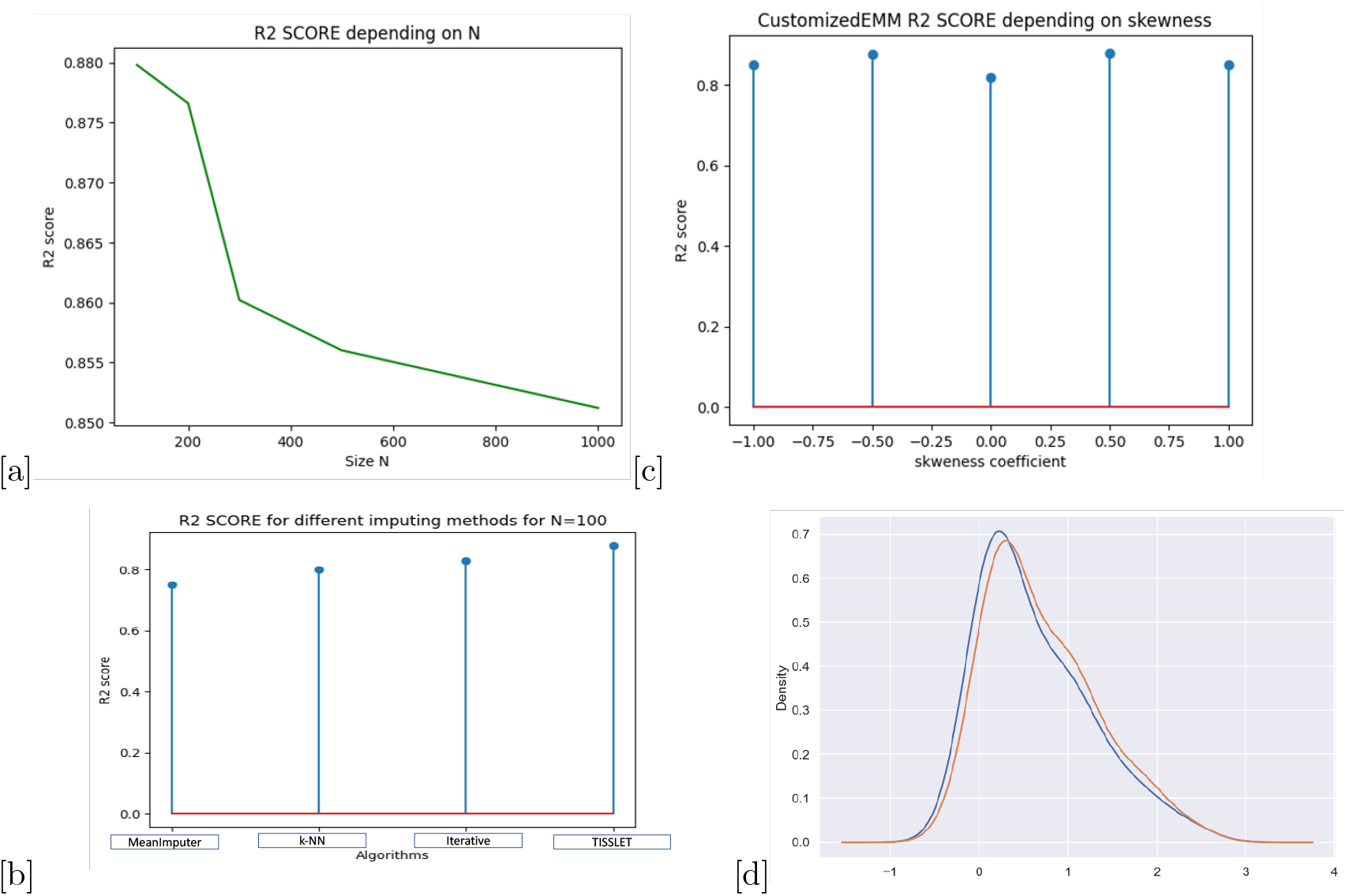
(a) Relationship between dataset size and R^2^ scores. TISSLET (CEM) R^2^ scores across skewness coefficients. (b) *A bar graph displaying the TISSLET framework’s R*^2^ *scores, indicating consistent predictive accuracy despite varying levels of skewness* {− 1.0; −0.5; 0; 0.5; 1.0} *in gene expression data*. (c) R^2^ score comparison among imputing methods for N=100. (d) *Bar chart illustrating R*^2^ *scores for different imputing methods, highlighting the superior performance of the TISSLET framework over MEANimputer, k-Nearest Neighbor (k=2), and iterative methods*.

Despite this, the TISSLET framework displayed a remarkable resilience in R^2^ scores across varying levels of skewness in gene expression data (Figure 5b), attesting to its robustness and adaptability to the diverse data distributions encountered in multiomics research. Further illustrating its efficacy, the TISSLET method demonstrated superior predictive accuracy when compared with other imputation methods such as MEANimputer, *k*-NN and iterative imputation for multi-tissue gene expression prediction (Figure 5c), reinforcing its suitability for complex genomic analyses.

#### Impact of the number of causal variants in simulation study

We extended our simulation study to assess the impact of a larger number of causal variants, ranging from 5 to 125, across a wide range of heritability levels. We found that increasing the number of causal variants had very little effect on the predictive performance of TISSLET. We believe this is because the expected imputation accuracy primarily depends on the total heritability explained by the causal SNPs (Figure 6).

**Fig. 6.**
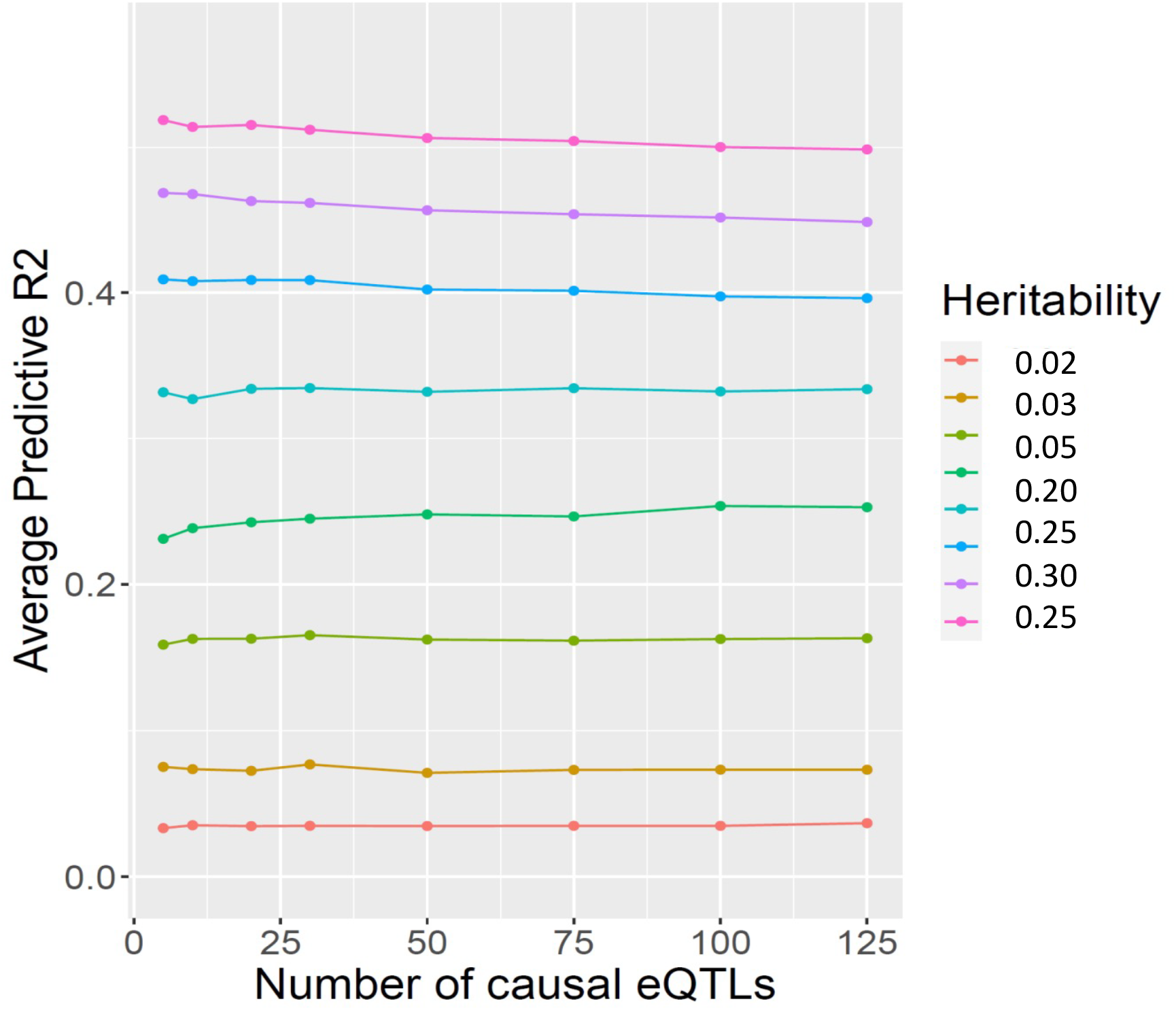
Impact of number of causal variants in simulation study.

#### Performance of fitting the skewed normal versus normal on predicted gene

Finally the density plot provides a visual comparison between the distributions of gene expression predictions using two [8] and our method of imputation. The curves are closely aligned and have a similar shape, peaking around a value of 1, which suggests that both methods yield relatively similar predictions in terms of their distribution. However, one curve is slightly shifted towards the right indicating that our approach may predict slightly higher expression values compared to the one that uses Normality assumption (Figure 5d).

#### Results of for SPATC1L gene study

In this section, we present results with tissue-tissue correlation for several errors, where *ρ* ∈ {0; 0.1; 0.3; 0.5; 0.7} varying. In this setting, we observe that our method, TISSLET performs better than all realistic competitors: only G-Lasso, outperforms slightly our method when *ρ* is less or equal than 0.015. As one would expect, when expression is nearly uncorrelated (*ρ* = 0), our method TISSLET performs better than all the others 3 methods that does implicitly assumes no tissue-tissue correlation. Remarkably, when *ρ* is greater than or equal to 0.3, TISSLET outperforms even the normal method which assume normality and missingness. In fact, the prediction accuracy of TISSLET increases as *ρ* increases, whereas all other methods, which do not explicitly model tissue-tissue correlation, have prediction accuracy remaining constant or slightly decreasing as *ρ* increases. This demonstrates the benefit of not only accounting for tissue-tissue correlation in multi-tissue joint eQTL mapping but also assume the correct distribution of the gene expression when expression across tissue types can be reasonably assumed to be conditionally dependent (Figure 7).

**Fig. 7.**
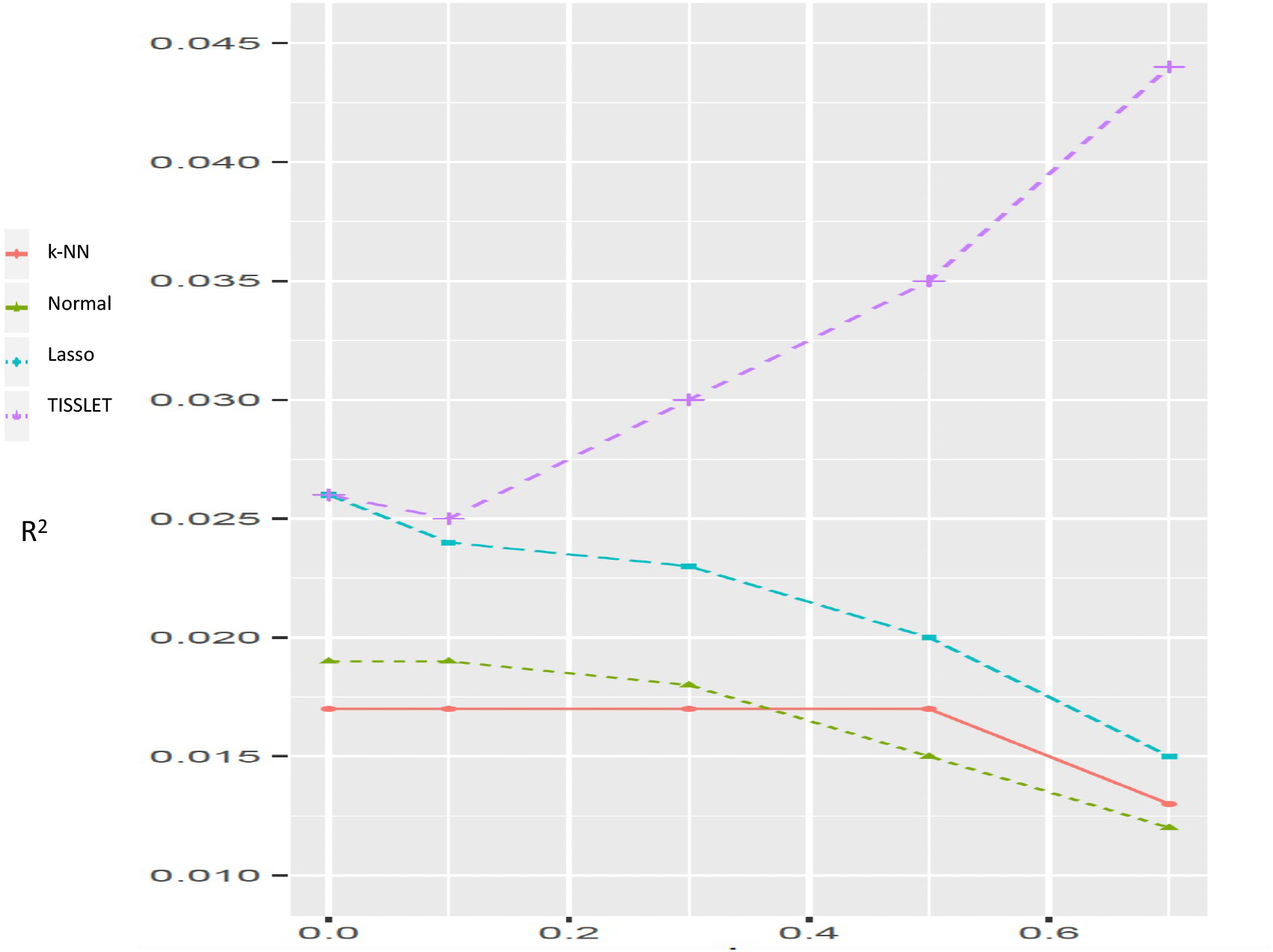
Average test R^2^ for 4 competing methods where *ρ* the correlation of the errors varies from 0.0 to 0.8.

Furthermore, the heatmap of gene expressions presented in Figure 8 illustrates a detailed example of a specific gene. In this case, out of the eight eQTLs (expression quantitative trait loci) that have been identified for this gene, seven of these eQTLs are consistently shared across all 29 examined tissues, demonstrating a uniform regulatory effect. Additionally, one of the eQTLs is shared across 28 of the 29 tissues, indicating a high level of commonality with a single tissue exception. This comprehensive visualization underscores the widespread influence of these eQTLs on gene expression across a broad range of tissue types, highlighting their significant role in the regulatory network.

**Fig. 8.**
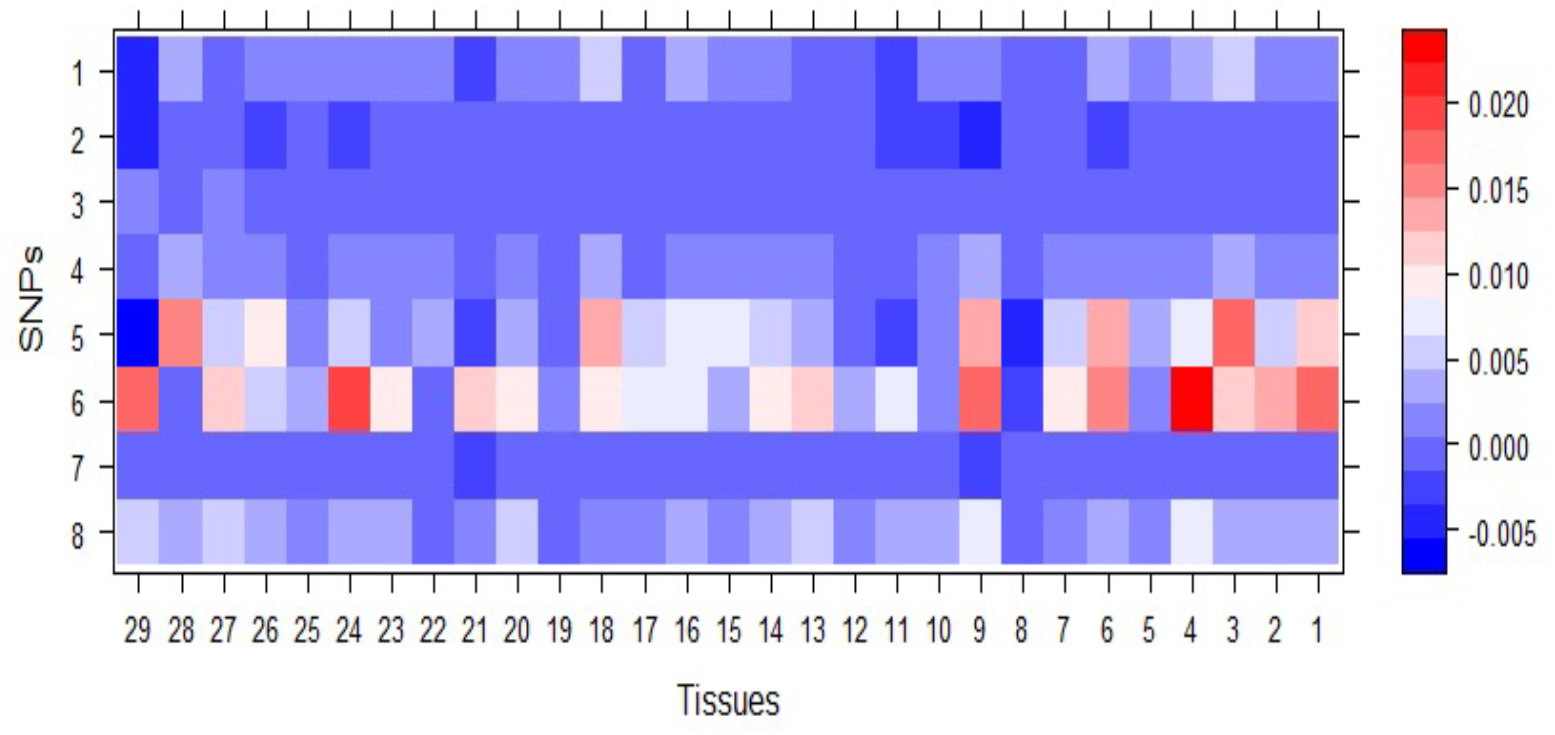
A heatmap depicting the expression of SNPs per tissue, with darker blue shades indicating weaker relationships

## 4 Conclusion and discussion

Our results suggest that the proposed TISSLET framework can robustly handle missing data in multi-tissue gene expression studies. The framework’s ability to retain accuracy across different imputation parameters, such as ‘k’, and its superior performance compared to other imputation methods, underscores its potential in improving complex trait predictions in multi-omics studies. The consistency of R^2^ scores regardless of skewness and dataset size provides evidence of the model’s stability. However, the observed decline in prediction accuracy with increasing dataset size suggests that further optimization of the framework may be necessary to manage the complexities of large-scale multi-omics data.

In conclusion, our findings advocate for the TISSLET framework’s application in multi-omics studies to improve the prediction of complex traits. Its ability to accurately account for tissue-tissue correlations and withstand data distribution asymmetries positions it as a powerful tool in the advancement of personalized medicine. As the complexity of multi-omics studies grows, the development of robust statistical machine learning like TISSLET is crucial for unveiling the genetic foundations of complex traits and advancing our understanding of gene expression dynamics.

While the TISSLET framework marks a significant advancement in gene expression prediction across multiple tissues, this study demonstrates its precision and stability, showcasing its high accuracy in predictions. However, the framework is still in its foundational stages and requires further refinement. The next steps involve optimizing the model to enhance its performance, particularly for large-scale datasets, thereby fully realizing its potential in complex multi-omics research and personalized medicine.

## 5 Appendix

### 5.1 Definition and notation

- 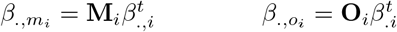
- 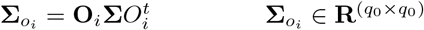
- 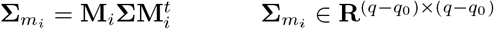
- 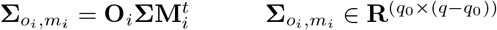
- 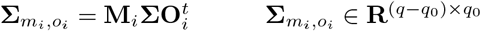
- 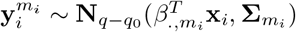
- 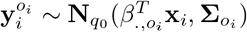

### 5.2 Penalized normal log-likelihood

Using Aaron et al. (2020), then can summarize the derivation of EM for estimating missing gene expression *ŷ*_*i*_, eQTL weights *β* and cross-tissue correlation Ω.

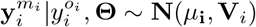

where

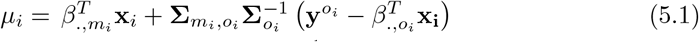

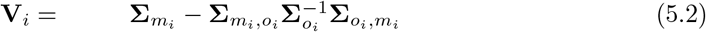

Then the negative log-likelihood of the **conditional** multivariate normal distribution for 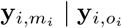 is proportional to

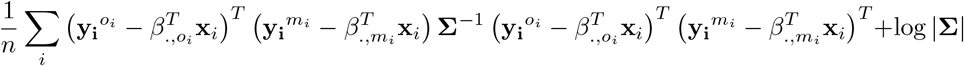

The negative log likelihood is useful in the EM steps because we needed to calculate the expectation of the log-likelihood *Q*(Θ):

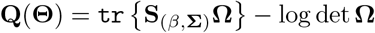

We regularize the entries of *β* and **Ω** using *l*_1_ penalties, and estimate them by minimizing the penalties likelihood

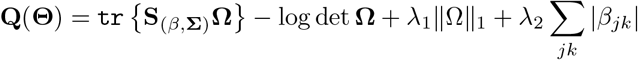

where the empirical covariance matrix

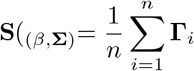

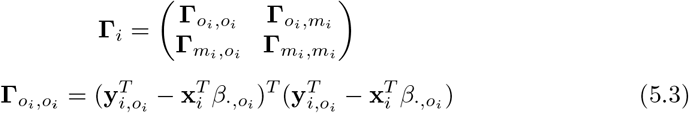

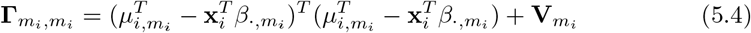

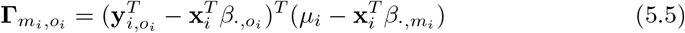

Then the algorithm is summarized as the following:

### 5.3 Penalized skew-normal log-likelihood

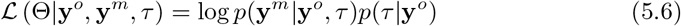

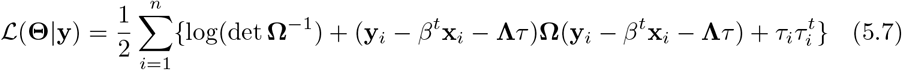

where **Θ** = (*β*, **Ω, Λ**) represents all unknown parameters. In the E-step of ECM, we need to calculate the Q-function, which is the conditional expectation of the complete data log-likelihood function (7) given the observed data **y**^*o*^ and the current estimate 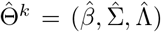. Herein, the term 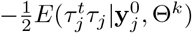 can be omitted because it does not include any parameters. Therefore, we have

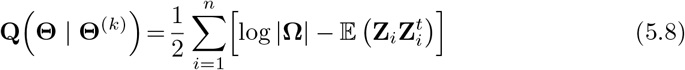

Using the formula 𝔼 (**ZZ**^*t*^) = Var(**Z**) + 𝔼 (**Z**) 𝔼 (**Z**)^*t*^, we have:

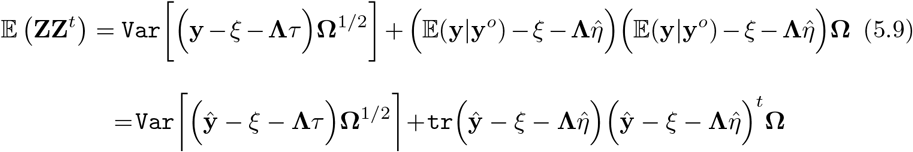

Using iterative expectation and 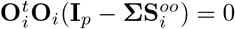, we have

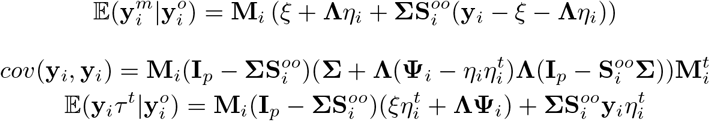

After simplification we have

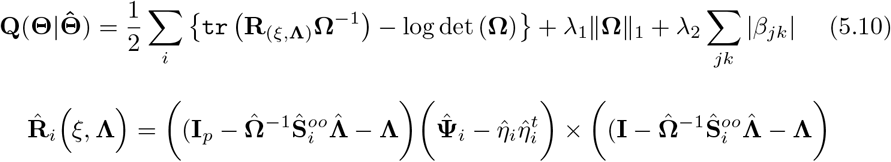

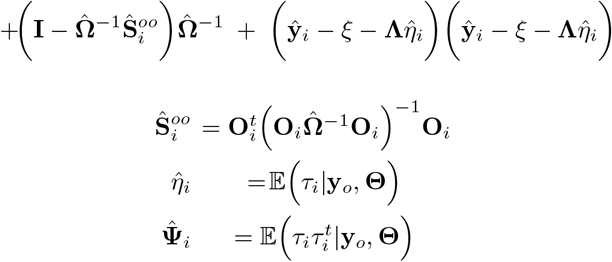

The predicted gene expression **ŷ**_*i*_ is given by

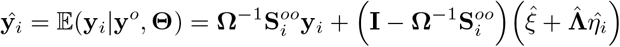

Using those derivation, Algorithm becomes:

#### Algorithm 3

Regularized EM Algorithm with skewed-normality.

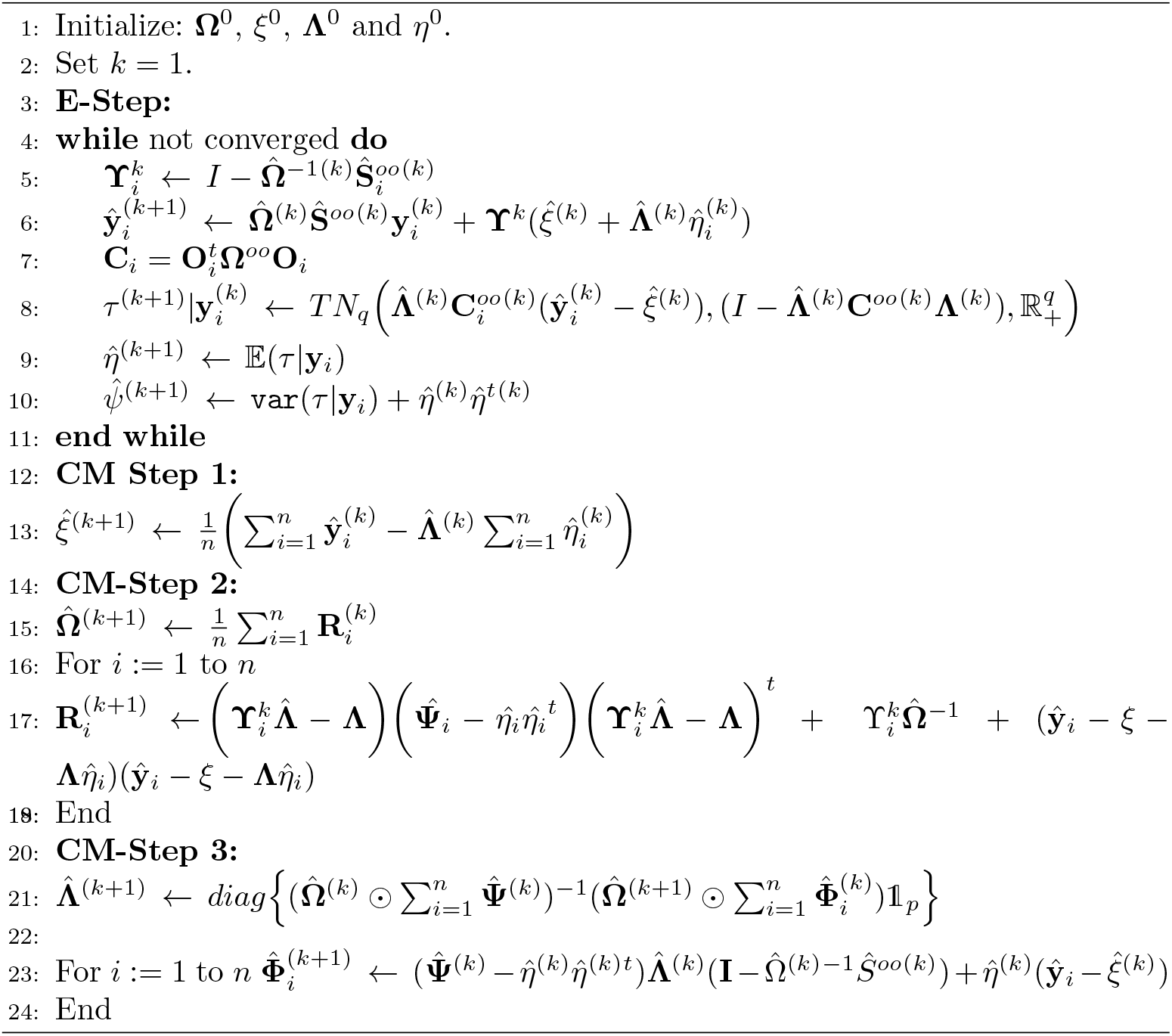

### 5.4 Precision matrix estimator

To estimate the inverse of the covariance matrix, several thresholding approaches have been proposed. The two most popular ones that we propose to use in this paper are GLASSO, available in R, that was proposed par Friedman et al. (2007) [16] and CLIME that was proposed by Cai et al. (2011) [18] If we use CLIME from Cai and Zhou approach, their equation emphasizes the fact that the solution may not be unique. Such nonuniqueness always occurs sinceΣ= **XX**^*T*^ where **X** ∈ ℝ^*n×p*^ is the sample data matrix and rank(**X**) ≤*n* ≪ *p* which is our case. However, there is no guarantee that the thresholding estimator is always positive definite [19]. Although the positive definite property is guaranteed in the asymptotic setting with high probability, the actual estimator can be an indefinite matrix, especially in real data analysis.

In the following we will describe quickly the argument and parameters of implementing BigQuiC in R.

### 5.5 Previous ad-hoc process for association SNPs with tissues

1. Create a copy of the genotype data matrix for each tissue. Only keep the samples that are matched with the corresponding gene expression and covariates data.
2. Remove SNPs with low minor allele frequency (MAF).
3. Remove genes with consistently low expression level.
4. For each tissue, load the tailored genotype data, gene expression data, covariates data, and gene/SNP location files into R.
5. To do a single tissue analysis, we will feed the data into MatrixEQTL to conduct single-tissue eQTL analysis and obtain summary statistics (i.e., t-statistics).
6. Using the cis window size to be 1 kb (i.e., 1 × 105) or 1 Mb (i.e., 1 × 106) and the p value threshold to be 1 in order to output summary statistics for all cis gene-SNP pairs then we convert each summary statistic *t* to a correlation *r* in each tissue 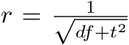 where df is the degree of freedom (the number of samples minus the number of covariates minus two) in the corresponding tissue.
7. Further we convert the correlations to z-statistics using Fisher transformation *z* = 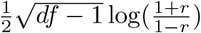.
8. Then we get the list of common gene-SNP pairs across all tissues and curate the obtained z-statistics into a matrix where each row corresponds to a gene-SNP pair in the common list and each column corresponds to a tissue. The z-statistics matrix is all we need for subsequent analyses.

## Author Contributions

The mathematical model was developed by H.B., writing the theoretical part and overview revision of the paper. M.C. Revised the paper. The author A.M. worked on running the mathematical model and producing the results. The author T.H. worked on producing the results in the earlier stage. The author A.A. worked on writing the results and discussion part with H.B. and A.M.; The corresponding author for submitting the article is A.A..

Declaration and Acknowledgements

We extend our gratitude to Dr. Molstad from the University of Minnesota for his invaluable assistance in providing access to his code and offering detailed explanations.

## Abbreviations

Not Applicable.

## Ethics approval and consent to participate

Not Applicable.

## Availability of data and material

Not Applicable.

